# RSPO/LGR signaling mediates astrocyte-induced proliferation of adult hippocampal neural stem cells

**DOI:** 10.1101/2023.12.18.572207

**Authors:** Daniela Valenzuela-Bezanilla, Muriel D. Mardones, Maximiliano Galassi, Sebastian B. Arredondo, Sebastian H. Santibanez, Nicolás Merino-Véliz, Fernando J. Bustos, Lorena Varela-Nallar

## Abstract

In the dentate gyrus of the adult hippocampus, neurogenesis from neural stem cells (NSCs) is regulated by Wnt signals from the local microenvironment. The Wnt/β-catenin pathway is active in NSCs, where it regulates proliferation and fate commitment, and subsequently its activity is strongly attenuated. The mechanisms controlling this pattern of activity are poorly understood. In stem cells from adult peripheral tissues, secreted R-spondin proteins (RSPO1-4) interact with LGR4-6 receptors and control Wnt signaling strength. Here, we found that RSPO1-3 and LGR4-6 are expressed in the adult dentate gyrus and in cultured NSCs isolated from the adult mouse hippocampus. The expression of LGR4-5 decreased in NSCs upon differentiation, concomitantly with the reported decrease in Wnt activity. Treatment with RSPO1-3 increased hippocampal NSCs proliferation and the expression of the Wnt target gene Cyclin D1. Moreover, RSPO1-3 were expressed by primary cultures of dentate gyrus astrocytes, a crucial component of the neurogenic niche able to induce NSC proliferation and neurogenesis. In co-culture experiments, astrocyte-induced proliferation of NSCs was prevented by RSPO2 knockdown in astrocytes, and by LGR5 knockdown in hippocampal NSCs. Altogether, our results indicate that RSPO/LGR signaling is present in the dentate niche, where it could control Wnt activity and proliferation of NSCs.

## INTRODUCTION

In the adult brain, neural stem cells (NSCs) are present in the subgranular zone (SGZ) of the hippocampal dentate gyrus. NSCs proliferate and differentiate into excitatory glutamatergic granule neurons that integrate into the dentate gyrus circuit and contribute to hippocampal function, a process termed adult hippocampal neurogenesis (Anacker and Hen 2017; Toda and Gage 2018; Kempermann 2022). The generation of functional adult-born granule neurons implies a fine-tuned regulation of the multiple steps of neurogenesis, including proliferation, fate commitment, differentiation, and maturation. NSCs in the SGZ are mostly quiescent (Shin et al. 2015; Urban and Cheung 2021), and once activated, they divide to generate actively proliferating neural progenitor cells (NPCs) that become neuroblasts, which differentiate into immature neurons and then into mature granule cells (Bonaguidi et al. 2011; Encinas et al. 2011).

The stages of neurogenesis are regulated by signaling factors secreted by different cell types found in the dentate niche. Among these factors, Wnt proteins play multiple roles during the process of neurogenesis [reviewed in (Arredondo et al. 2020b)]. Moreover, astrocytes that are major components of the adult dentate gyrus (Rieskamp et al. 2022), are known to regulate the maintenance of adult NSCs and induce neurogenesis in a Wnt-dependent manner (Lie et al. 2005; Wexler et al. 2009; Okamoto et al. 2011). Wnts are secreted glycoproteins that bind to seven-pass transmembrane Frizzled (FZD) receptors, which can trigger the canonical Wnt/β-catenin signaling pathway, or the non-canonical (β-catenin-independent) signaling cascades (Kohn and Moon 2005; Gordon and Nusse 2006; Butler and Wallingford 2017). Activation of the canonical Wnt/β-catenin signaling involves the formation of a ternary complex composed of Wnt, FZD, and the co-receptor low density lipoprotein receptor-related protein 5 or 6 (LRP5/6). Wnt stimulation results in the stabilization and nuclear import of β-catenin, which acts as a transcriptional co-activator interacting with members of the T cell factor/lymphoid enhancer binding factor (TCF/LEF) family of transcription factors to induce the expression of Wnt target genes (e.g., CyclinD1 and Axin2 (Shtutman et al. 1999; Jho et al. 2002; Lustig et al. 2002)). The Wnt/β-catenin pathway controls cell fate decisions during embryonic development and in adult organisms, acting as a stem cell niche signal in many tissues (Gordon and Nusse 2006; Nusse and Clevers 2017). In adult neurogenesis, Wnt/β-catenin signaling controls proliferation and fate commitment of neural precursor cells (Kuwabara et al. 2009; Mao et al. 2009; Wexler et al. 2009; Qu et al. 2013; Austin et al. 2021), and in maturing newborn neurons, it regulates dendritic arbor complexity (Heppt et al. 2020). The role of the Wnt proteins has been studied in cultured NSCs isolated from the adult mouse hippocampus. These cells can differentiate into neurons or glial cells (Babu et al. 2011), and represent a controlled *in vitro* system to evaluate neurogenesis. In hippocampal NSCs, Wnt proteins induce cell proliferation through the expression of CyclinD1 (Wexler et al. 2009; Qu et al. 2013; Austin et al. 2021) and fate commitment through the expression of Neurogenin 2 (Ngn2) and NeuroD1 (Kuwabara et al. 2009; Qu et al. 2013; Amador-Arjona et al. 2015).

Wnt/β-catenin reporter mice have shown that this signaling pathway is active in NSCs, but then its activity is strongly attenuated (Lie et al. 2005; Garbe and Ring 2012; Heppt et al. 2020; Austin et al. 2021), which was proposed to be important for the proper dendritogenesis of newborn neurons (Heppt et al. 2020). The mechanisms involved in the dynamics of Wnt/β-catenin signaling during the early stages of adult neurogenesis are poorly understood. During development and in adult peripheral tissues, the strength of the Wnt/β-catenin signaling pathway is tightly regulated by the roof plate-specific spondin (R-spondin, RSPO) proteins (de Lau et al. 2012). RSPOs are a family of four cysteine-rich secreted glycoproteins (RSPO1-4) that interact with the cell surface leucine-rich repeat-containing G protein-coupled receptors (LGR4-6) and with the transmembrane E3 ubiquitin ligases zinc and ring finger 3 (ZNRF3) or ring finger protein 43 (RNF43), which in the absence of RSPO ubiquitinate FZDs inducing their endocytosis and degradation (Carmon et al. 2011; de Lau et al. 2011; Glinka et al. 2011; Gong et al. 2012; Zebisch and Jones 2015). RSPO/LGR interaction prevents FZD ubiquitination, thereby augmenting Wnt/β-catenin signaling activity (Hao et al. 2012; Koo et al. 2012; Schuijers and Clevers 2012). Through this mechanism, RSPO proteins control proliferation and differentiation of stem cells in several adult tissues, including the intestine, stomach, hair follicles, and liver, among others (de Lau et al. 2014; Nagano 2019). Whether RSPO/LGR signaling plays a role in adult NSC behavior and adult neurogenesis is unknown. Here, we studied whether RSPO proteins and LGR receptors are expressed in the adult mouse dentate gyrus and evaluated whether RSPO proteins regulate proliferation of hippocampal NSCs. In addition, we evaluated the role of RSPO signaling in astrocyte-induced proliferation of neural precursors. Our results strongly suggest that RSPO/LGR is part of the signaling mechanisms present in the adult dentate niche that control the proliferation of NSCs.

## RESULTS

### RSPO proteins and LGR receptors are expressed in the adult mouse dentate gyrus

To investigate whether RSPO signaling exists in the dentate niche, we evaluated the expression of RSPO proteins and their LGR4-6 receptors in the dentate gyrus of 2-month-old mice. RSPO1-3 mRNA and protein levels were analyzed by RT-qPCR and immunoblotting (Fig. 1A-C). The variations in molecular weights, and the presence of more than one band are expected due to glycosylation of RSPO proteins. Structurally, these proteins comprise an N-terminal signal peptide, two cysteine-rich furin-like domains (Fu1 and Fu2), followed by a thrombospondin type 1 repeat domain (TSR-1), an a basic charged C-terminal tail (Kim et al. 2006; Zebisch et al. 2013). RSPO1 and RSPO3 are both N-glycosylated at N137 in the Fu2 domain, and RSPO2 is N-glycosylated at N160 in the TSR-1 domain, which are crucial for efficient folding for secession (Chang et al. 2016). LGR4-6 mRNAs were also detected in the adult mouse dentate gyrus (Fig. 1A, B), and LGR4-5 were assessed at the protein level (Fig. 1C).

**Figure 1.**
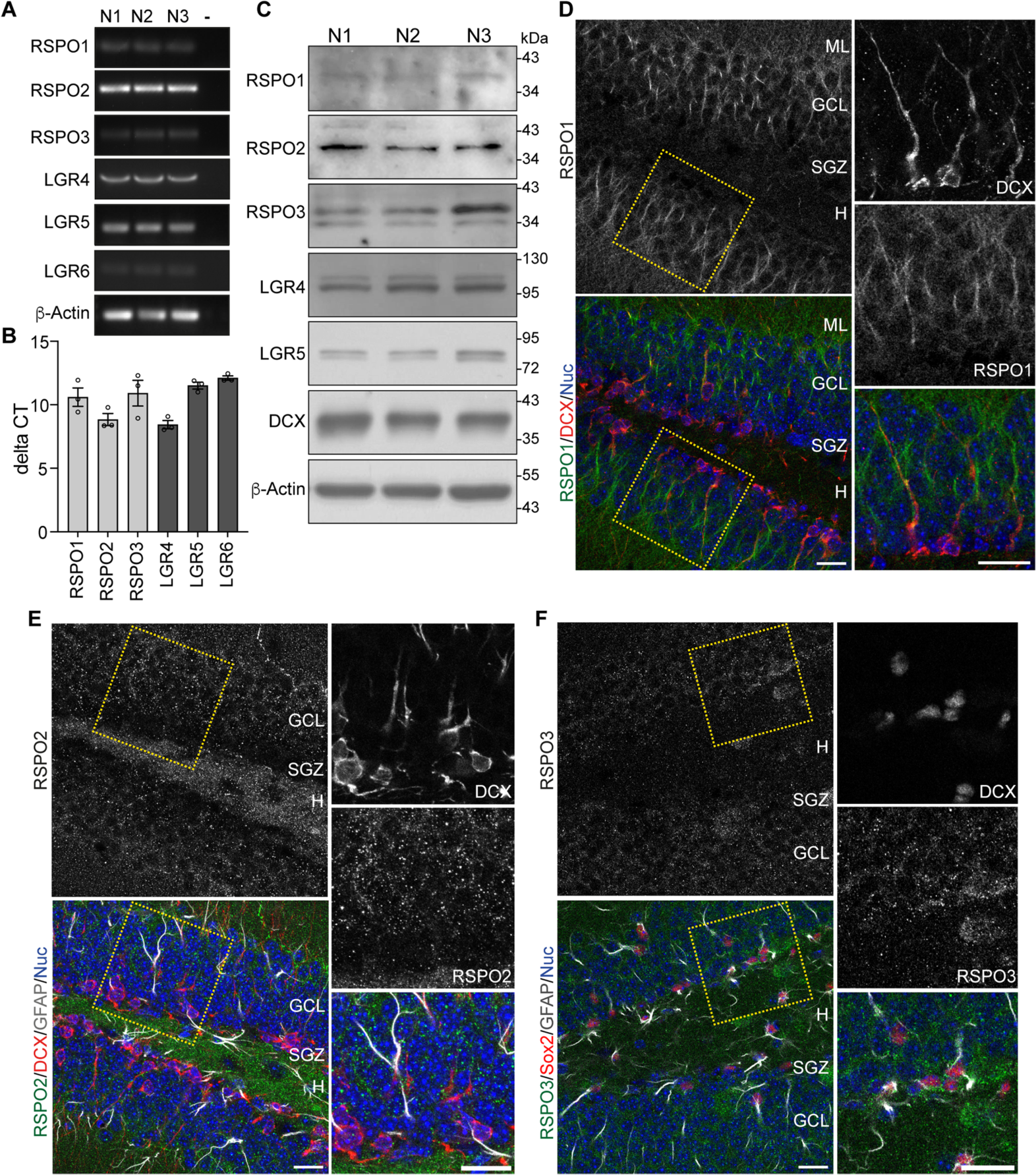
RSPO proteins and LGR receptors are expressed in the dentate gyrus of the adult mouse brain. (A, B) RT-qPCR from total RNA isolated from the dentate gyrus of 2-month-old mice. (A) End-point RT-qPCR reaction products. (B) delta CT values normalized to β-actin used as a reference gene. Of note, the lower the delta Ct value, the higher the expression of the gene. Bars show mean ± SEM. Dots show individual data (each one corresponds to 2 mice; total 6 mice). (C) Immunoblot from total protein extracts of the dentate gyrus of 2-month-old mice. Each N represents a pool of extracts from two animals. Numbers on the right indicate molecular weight standards (kDa). (D-F) Representative immunofluorescence staining of RSPO1 (C), RSPO2 (D), and RSPO3 (E) in the dentate gyrus of a 2-month-old mouse. Higher magnifications of the dotted squares are shown at the right. Scale bars: 20 μm. Immunodetection of DCX (red) was used as a marker of immature neurons in C and D; and GFAP (gray) and Sox2 (red) were used as markers of NSCs and astrocytes in E. NucBlue (Nuc, blue) was used for nuclear staining. GCL: granule cell layer; ML: molecular layer; SGZ: subgranular zone; H: Hilus.

The distribution of RSPO proteins in the dentate gyrus was investigated by immunofluorescence staining. RSPO1 was mainly detected in the middle and outer thirds of the granule cell layer (GCL) and in the molecular layer (Fig. 1D), suggesting RSPO1 is distributed close to the dendritic arbor of granule cells. In agreement, co-distribution of RSPO1 with dendrites of immature neurons positive for doublecortin (DCX) was found in the GCL (Fig. 1D). Even though there was also RSPO2 staining in the middle and outer thirds of the GCL, it was mainly detected in the hilus (Fig. 1E). RSPO3 staining showed a low and homogenous distribution in the GCL, molecular layer, and hilus (Fig. 1E), and was detected in the SGZ (Fig. 1E).

RSPOs staining was also evaluated in the CA3 region, where the axons of newborn neurons are projected (Zhao et al. 2006). RSPO1 staining showed a somatodendritic distribution in CA3 neurons; RSPO2 staining was mainly observed in the *stratum lucidum*, suggesting an axonal distribution in mossy fibers; and RSPO3 showed a very low staining in CA3 neurons (Supplemental Fig. S1).

Altogether, our data indicate that RSPO proteins and LGR receptors are expressed in the adult mouse dentate gyrus, suggesting that RSPO/LGR signaling occurs in the dentate niche.

### RSPO proteins induce proliferation of adult hippocampal NSCs

Considering that RSPO1-3 are expressed in the dentate gyrus, we evaluated whether these proteins regulate adult NSCs. For this experiment, we used cultured NSCs isolated from the adult mouse hippocampus as an *in vitro* model of neurogenesis (Babu et al. 2011; Arredondo et al. 2020a). We found that these cells express RSPO1-3 mRNA and protein (Fig. 2A-C). The expression of LGR4-6 mRNA was detected by RT-qPCR, and the results suggest that LGR6 exhibited the lowest level of expression in NSCs (Fig. 2A, B). LGR4-5 proteins were determined by immunoblotting (Fig. 2C) and immunostaining (Fig. 2D). LGR4-5 staining was detected in all cells positive for the neural stem/progenitor cell marker Nestin (Fig. 2C), indicating that hippocampal NSCs express the receptors for RSPO proteins. Interestingly, upon differentiation induced by growth factors withdrawal, LGR4-5 mRNA levels decreased significantly (Fig. 2E, F), while the mRNA of the neuronal marker Ngn2 increased (Fig. 2G). In addition, LGR5 protein levels significantly decreased, and LGR4 tended to decrease upon differentiation (Fig. 2H-K), which coincided with an increase in the immature neuronal marker DCX (Fig. 2H, I). These results indicate that the expression of LGR receptors is reduced upon differentiation, suggesting that RSPO/LGR signaling might have a role in proliferative NSCs.

**Figure 2.**
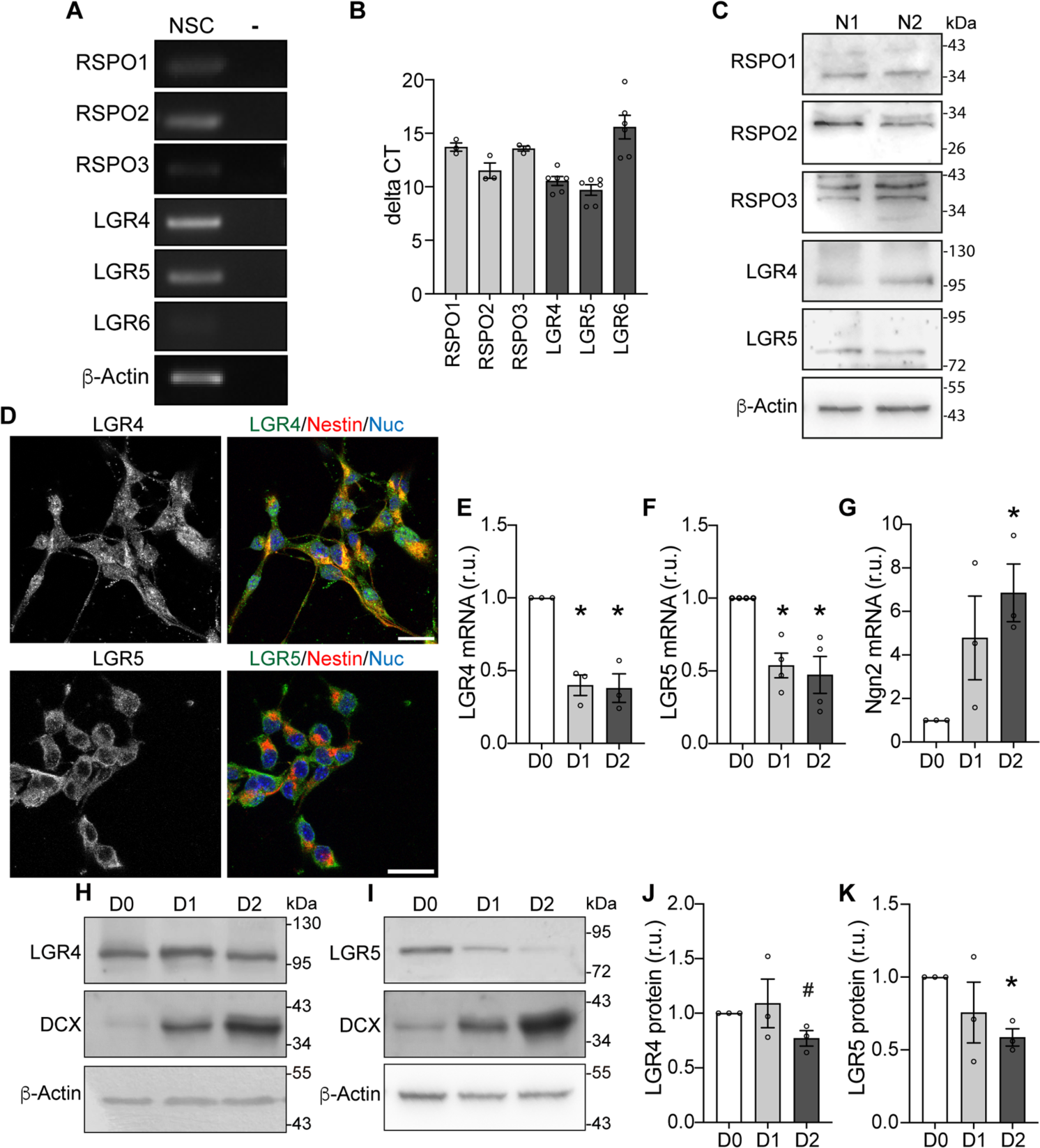
Expression of RSPO proteins and LGR receptors in cultured adult hippocampal NSCs. (A, B) RT-qPCR from total RNA isolated from cultured NSCs. End-point RT-qPCR reaction products (A) and delta CT values normalized to β-actin (B) are shown. Of note, the lower the delta Ct value, the higher the expression of the gene. Bars show mean ± SEM. (C) Immunoblot from total protein extracts of NSCs. The numbers on the right indicate the molecular weight standards (kDa). (D) Representative immunofluorescence staining of LGR4 or LGR5 (green), Nestin (red), and the nuclear staining NucBlue (Nuc, blue). Scale bar: 20 μm (E-G) RT-qPCR from total RNA isolated from NSCs under proliferative conditions (D0) and 24 (D1) or 48 (D2) hours after inducing differentiation by growth factor withdrawal. The mRNA levels of LGR4, LGR5, and Ngn2 (used as differentiation control) were normalized to β-actin mRNA and expressed relative to D0. (H-I) Immunoblot analysis of LGR4-5 in total protein extracts from NSCs at D0, D1, and D2. DCX was used as a neuronal differentiation control. (J-K) Densitometric analysis of LGR4-5 and DCX was normalized to β-actin and expressed relative to D0. Bars show mean ± SEM, and dots represent independent experiments. A one-sample t-test and Wilcoxon test were used. *p < 0.05, #p < 0.08.

To evaluate the effect of RSPOs on proliferation, NSCs were treated for 24 hours with 100 ng/ml recombinant RSPO1 (rRSPO1), rRSPO2, or rRSPO3 and incubated with the nucleotide analog BrdU for the last 2 hours of treatment. The incorporation of BrdU was analyzed by immunofluorescence staining (Fig. 3A). The percentage of BrdU positive cells was significantly increased in NSCs treated with rRSPO1-3 (Fig. 3B-D), indicating that treatment with RSPOs induced proliferation.

**Figure 3.**
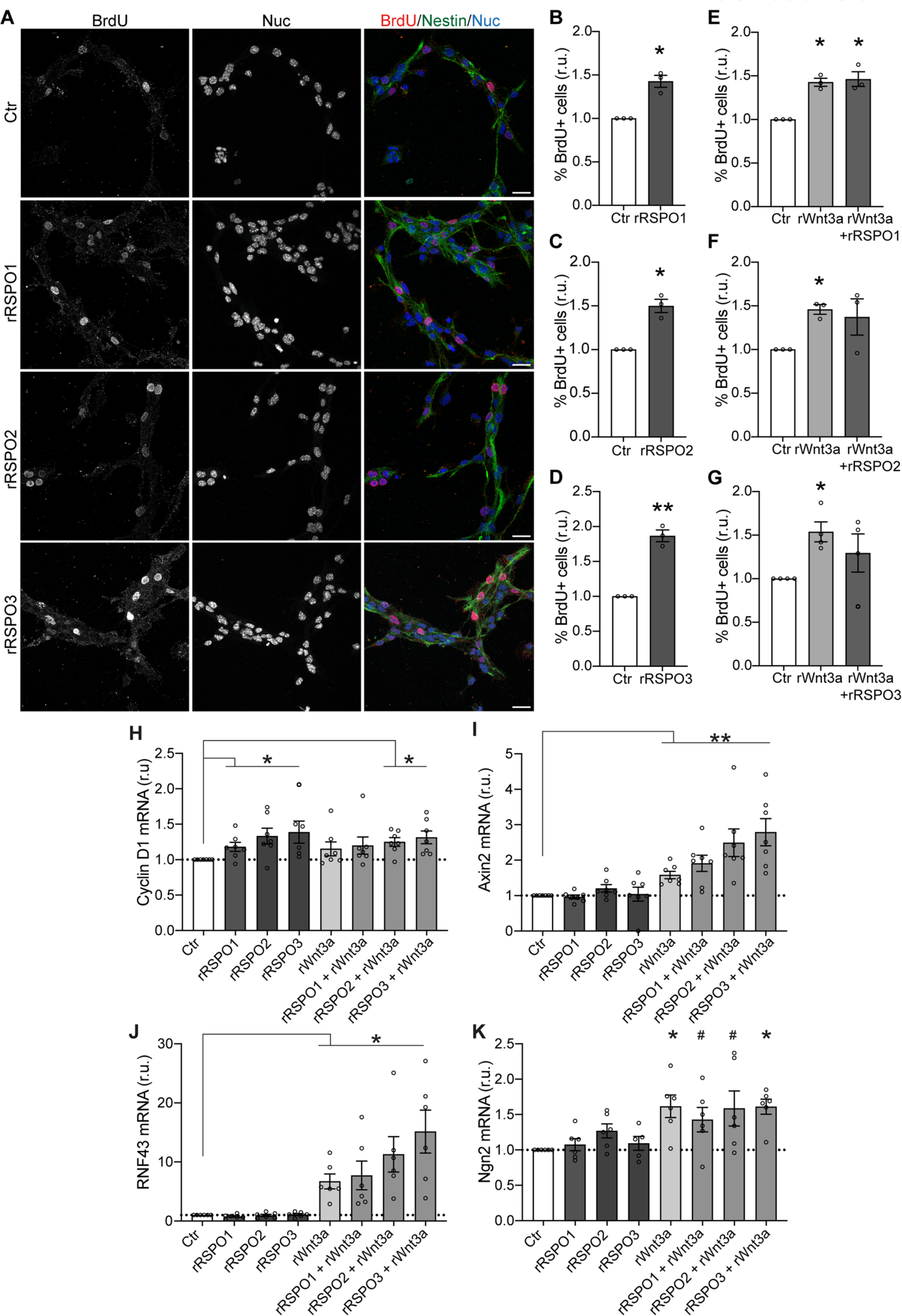
Treatment with RSPO proteins induces the proliferation of hippocampal NSCs and the expression of Wnt target genes. (A) Representative immunofluorescence staining of BrdU (red), Nestin (green) and Nuc (blue) in NSCs incubated for 24 hours in the presence or absence of 100 ng/ml recombinant RSPO1 (rRSPO1), rRSPO2, or rRSPO3 and were incubated with 10 μM BrdU for the last 2 hours of treatment. Scale bar: 20 μm. (B-G) Quantification of the percentage of BrdU positive NSCs treated with 100 ng/ml rRSPO1-3 (B-D) and 150 ng/ml of rWnt3a ± 100 ng/ml of rRSPO1-3 (E-G). Data are presented as fold-change relative to the control condition. Representative immunostaining of NSCs treated with rWnt3a ± rRSPO1-3 are shown in Supplemental Fig. 2. (H-K) RT-qPCR from total RNA isolated from NSCs treated for 24 hours with 100 ng/ml rRSPO1-3 or 150 ng/ml rWnt3a ± 100 ng/ml rRSPO1-3. Bars show mean ± SEM, and dots represent independent experiments. A one-sample t test and Wilcoxon test were used to compare all conditions versus the control condition. Kruskal-Wallis test was used for multiple comparisons. *p<0.05, **p<0.01, ***p<0.001, #p<0.06.

RSPOs potentiate Wnt signaling activity by augmenting the sensitivity of cells to Wnt ligands (Schuijers and Clevers 2012). Therefore, the effect of rRSPOs likely involves the enhancement of endogenous Wnt ligands action (Wexler et al. 2009). Intriguingly, we found that co-treatment with rRSPO1-3 plus recombinant Wnt3a (rWnt3a) ligand, known to induce proliferation of hippocampal NSCs (Wexler et al. 2009), did not augment the effect of rWnt3a (Fig. 3E-G and Supplemental Fig. S2). This result indicates that RSPOs were not able to augment the effect of an exogenous Wnt on NSCs proliferation.

To evaluate whether treatment with rRSPOs alone or in combination with rWnt3a induced Wnt/β-catenin signaling activity, the expression of Wnt target genes was assessed by RT-qPCR. For this experiment, NSCs were treated with 100 ng/ml rRSPOs in the presence or absence of 150 ng/ml rWnt3a for 24 hours (Fig. 3H-K). We evaluated the expression of CyclinD1, previously shown to mediate the proliferative effect of Wnt/β-catenin signaling on cultured hippocampal NSCs (Qu et al. 2013). We found that treatment with rRSPO1-3 significantly increased CyclinD1 mRNA levels (Fig. 3H) but did not increase the effect of rWnt3a, in agreement with the finding that co-treatment did not potentiate the effect on proliferation (Fig. 3E-G). Interestingly, the expression of Axin2, a well-described Wnt target gene (Jho et al. 2002; Leung et al. 2002), was not induced by treatment with rRSPO1-3, while treatment with rWnt3a alone or in the presence of rRSPOs significantly increased Axin2 mRNA levels (Fig. 3I). Axin2 is a component of the complex that induces the degradation of β-catenin and, therefore, is a target gene that functions in a negative feedback loop to limit Wnt signaling activation (Jho et al. 2002; Lustig et al. 2002). Furthermore, treatment with rWnt3a alone or with rRSPOs strongly induced the expression of RNF43 (Fig. 3J), which also induces a negative feedback loop for Wnt activity through ubiquitination of FZD receptors (Koo et al. 2012). These results suggest that treatment with rWnt3a ± rRSPOs induces a negative regulation of the Wnt/β-catenin signaling, which is not induced by treatment with rRSPOs alone.

We also found that co-treatment with rWnt3a ± rRSPO increased Ngn2 mRNA (Fig. 3K). Ngn2 is a target gene involved in Wnt-mediated neuronal differentiation of hippocampal NSCs (Qu et al. 2013; Amador-Arjona et al. 2015). Interestingly, previous reports have shown that high stimulation of Wnt/β-catenin signaling in proliferating NSCs reduces their proliferative capacity and promotes neuronal differentiation (Austin et al. 2021). In this context, the lack of potentiation on proliferation and CyclinD1 expression by co-treatment with rRSPOs plus rWnt3a could be attributed to an increase in differentiation induced by higher Wnt activity.

### RSPO proteins mediate astrocyte-induced proliferation of adult hippocampal neural progenitor cells

Astrocytes are a crucial component of the dentate niche. *In vitro*, hippocampal astrocytes induce proliferation and neurogenesis of cultured NSCs (Song et al. 2002; Lie et al. 2005; Okamoto et al. 2011). To assess the potential role of RSPOs in the astrocyte-mediated induction of proliferation, we created sandwich co-cultures of NSCs with hippocampal astrocytes (NSC/Astro co-cultures), using a custom-made 3D-printed support (Supplemental Fig. S3A), in which cells shared the medium without direct contact with each other (Kaech and Banker 2006). For these experiments, astrocytes were isolated from the dentate gyrus of adult mice. As expected, these cells expressed the astrocytic markers GFAP and S100β (Supplemental Fig. S3B), and induced proliferation of NSCs in co-culture experiments (Supplemental Fig. S3C).

We found that astrocytes extracted from the dentate gyrus expressed RSPO1-3 mRNAs (Fig. 4A, B) and proteins (Fig. 4C). To investigate the contribution of RSPO proteins in astrocyte-induced proliferation of hippocampal NSCs, NSCs/Astro co-cultures were performed for 24 hours in the presence or absence of soluble Fc-LGR4 or Fc-LGR5 chimeras consisting of the extracellular domain of LGR receptors fused to the Fc region of immunoglobulin (Fig. 4D). Soluble Fc-LGRs bind to RSPO proteins, preventing their action. Proliferation of NSCs co-cultured with astrocytes was significantly reduced in the presence of Fc-LGR4 or Fc-LGR5 but was not affected by the control Fc protein (Fc-Ctrl) (Fig. 4E and Supplemental Fig. S4A). This result suggests that RSPO proteins are involved in the astrocyte-induced proliferation of NSCs. Noteworthy, when NSCs were cultured alone (not in co-cultures) and incubated with Fc-LGRs, there was no effect on proliferation (Supplemental Fig. S4B), indicating that the decrease in proliferation induced by the treatment with Fc-LGR4-5 in NSC/Astro co-cultures was mediated by RSPO ligands secreted by astrocytes and not by NSC-derived RSPOs.

**Figure 4.**
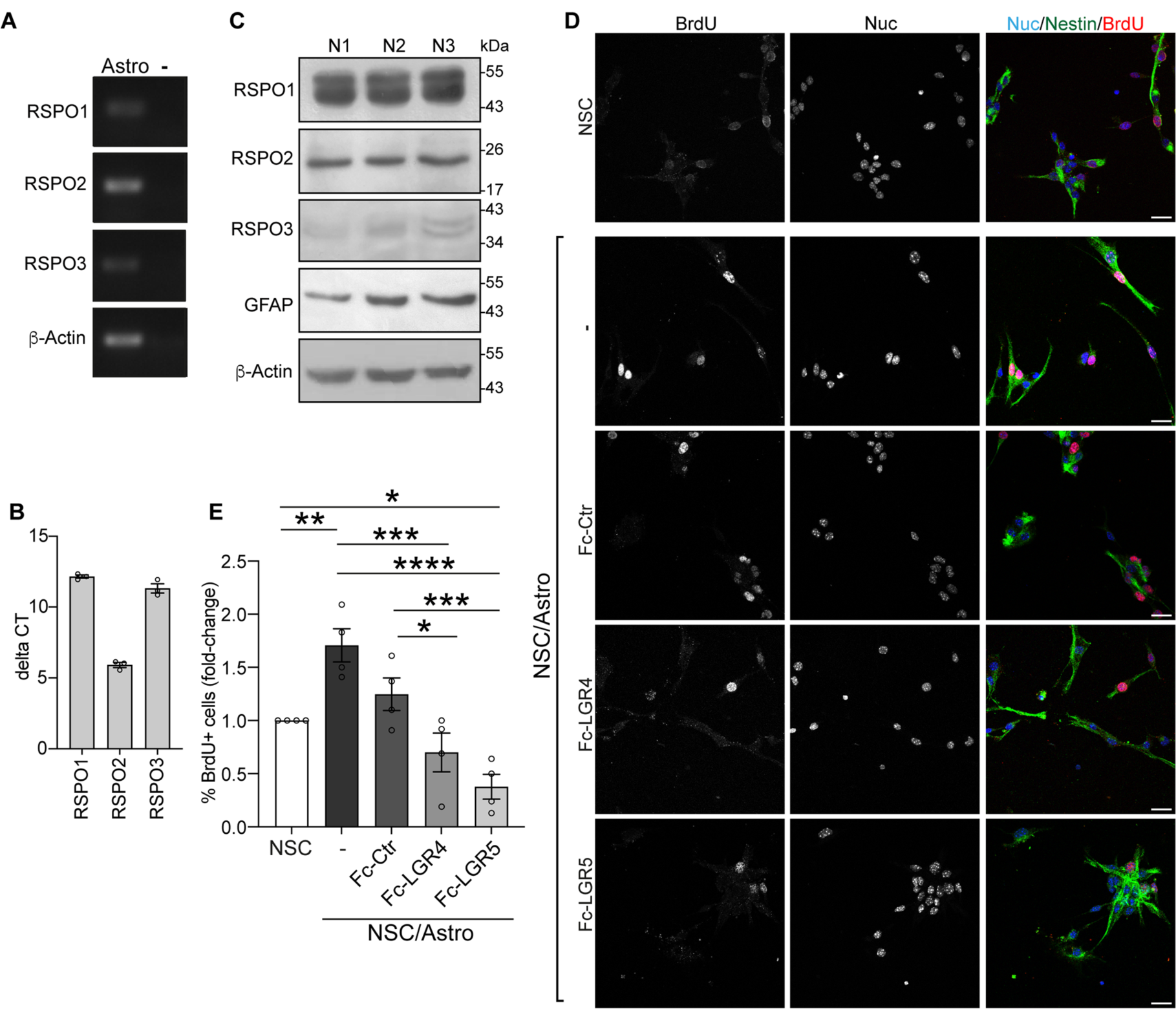
Cultured dentate gyrus astrocytes express RSPO proteins and induce proliferation of adult hippocampal NSCs in an RSPO-dependent manner. (A, B) RT-qPCR from total RNA isolated from primary cultured astrocytes isolated from the dentate gyrus of adult mice. End-point RT-qPCR reaction products (A) and delta CT values normalized to β-actin (B) are shown. Bars show mean ± SEM. (C) Immunoblot of total protein extracts from dentate gyrus astrocytes. Extracts from three different animals (N1–N3) are shown. The numbers on the right indicate the molecular weight standards (kDa). (D) Representative immunodetection of BrdU (red), Nestin (green), and Nuc (blue) in NSCs that were co-cultured (NSC/Astro) or not (NSC) with astrocytes for 24 hours in the presence or absence of 0.25 μg/ml Fc-LGR4, Fc-LGR5, or a control Fc (Fc-Ctrl) chimera, and BrdU was incorporated for the last 4 hours before fixation. Scale bar: 20 μm. (E) Percentage of cells positive for BrdU. Data are presented as fold-change relative to NSCs not co-cultured with astrocytes. Bars show mean ± SEM, and dots represent independent experiments. Two-way ANOVA followed by a Tukey’s multiple comparison test was used. *p<0.05, **p<0.01, ***p<0.001, and ****p<0.0001.

To further demonstrate the role of astrocyte-derived RSPOs on NSCs proliferation, we knocked down the expression of RSPO2, which is the most highly expressed in astrocytes (Fig. 4A, B). Knockdown was first assessed in Neuro2a (N2a) cells transduced with lentiviruses expressing two different shRNAs targeting RSPO2 (shR1 and shR2), or a control shRNA (shC), plus GFP. Both shR1 and shR2 significantly reduced RSPO2 protein levels compared with shC (Fig. 5A, B), and shR2 was selected for further experiments. In N2a cells, shR2 significantly reduced RSPO2 mRNA compared to control cells transduced with shC (Fig. 5C). Then, RSPO2 knockdown was assessed in astrocytes transfected with shC or shR2, and RSPO2 protein levels were analyzed in total cell lysates and in the secreted media (Fig. 5D), since RSPO2 is a secreted glycoprotein. Remarkably, the immunoreactive bands observed in the media displayed a reduced molecular mass than the bands observed in the lysates (Fig. 5D). In both, cell lysates (Fig. 5E) and media (Fig. 5F), reduced RSPO2 levels were observed in astrocytes transfected with shR2, compared to shC (Fig. 5E, F).

**Figure 5.**
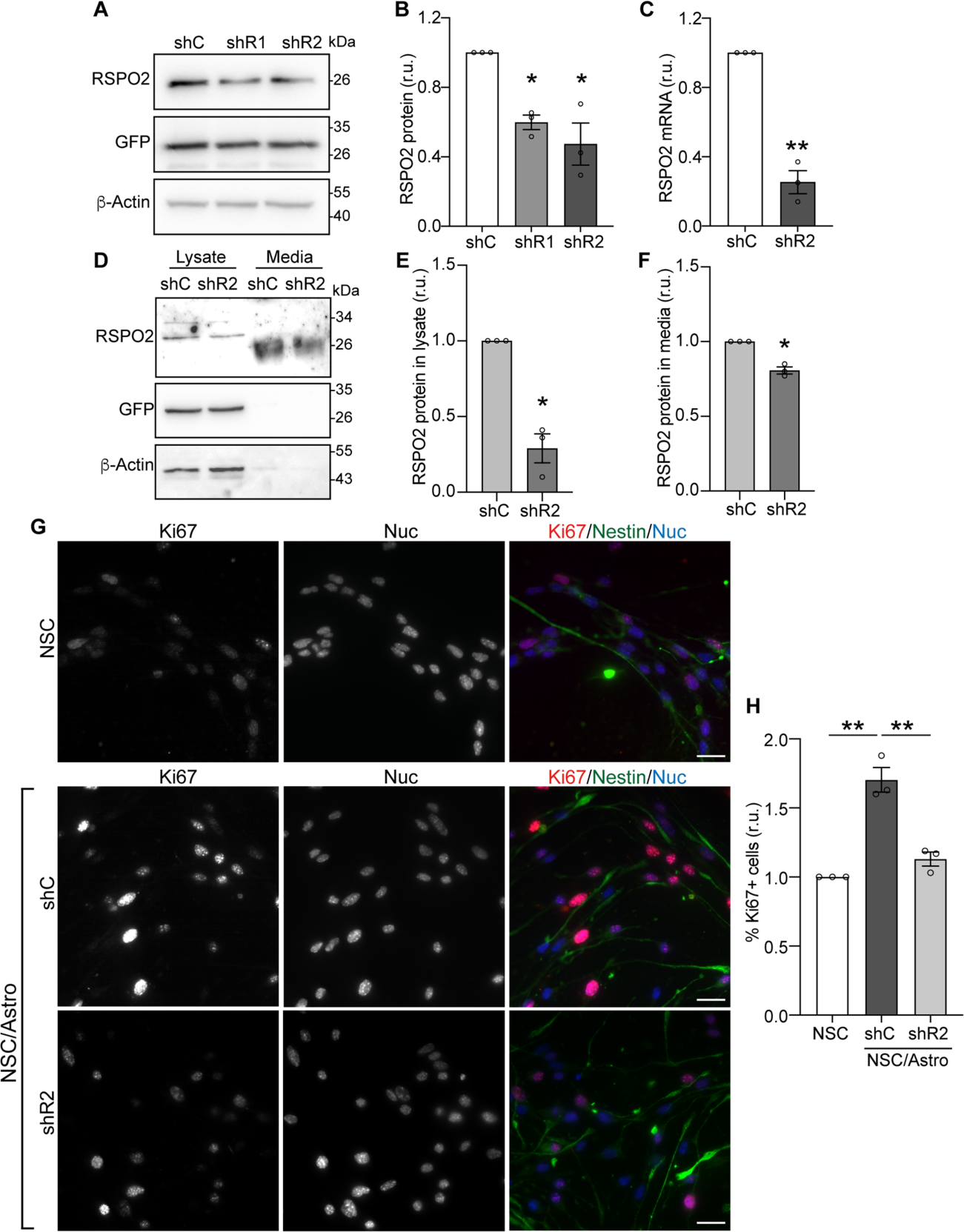
RSPO2 knockodown reduces astrocyte capacity to induce the proliferation of hippocampal NSCs. (A) Immunoblot of RSPO2 in the total protein extract of N2a cells transduced with lentivirus expressing shRNAs targeting RSPO2 (shR1, shR2) or a control shRNA (shC) and the reporter protein GFP. (B) Densitometric analysis of the immunoblots. RSPO2 protein levels were normalized to β-actin and expressed relative to shC. (C) RT-qPCR from total RNA isolated from N2a cells transduced for 72 hours with shR2 or shC. RSPO2 mRNA was normalized to β-actin mRNA and expressed relative to shC. (D) Immunoblot in total protein extracts (lysates) or the media of dentate gyrus astrocytes 72 hours after transfection with shR2 or shC vectors. (E-F) Densitometric analysis of the immunoblots. RSPO2 protein levels in the lysate were normalized to β-actin. RSPO2 levels in the lysates and media are expressed relative to the shC condition. (G) Representative immunodetection of Ki67 and Nestin in NSCs with or without co-culture for 24 hours with astrocytes transfected with shC or shR2. NucBlue (Nuc) was used as a nuclear marker. (H) Percentage of Ki67+ cells. Bars represent the mean ± SEM. Dots represent independent experiments. One-sample t-test and Wilcoxon test (B, C, E, F), or Two-way ANOVA followed by Tukey’s multiple comparison test (H) were used. *p<0.05, **p<0.01.

To evaluate the role of RSPO2 in astrocyte-induced proliferation, NSCs were co-cultured for 24 hours with astrocytes transfected with shC or shR2, and proliferation was assessed by the expression of the mitotic marker Ki67 (Fig. 5G). Compared to NSCs cultured without astrocytes, co-culture with shC-expressing astrocytes significantly increased the percentage of NSCs positive for Ki67, an effect that was significantly decreased in NSCs co-cultured with shR2-expressing astrocytes (Fig. 5H). This result indicates that RSPO2 is required for proliferation induced by astrocytes.

To further assess the role of RSPO/LGR signaling in astrocyte-induced proliferation, LGR5 was knocked down in NSCs by transduction with a retrovirus expressing an shRNA targeting LGR5 (shLGR5). The retroviruses expressing shLGR5 or a control shRNA (shC) also expressed the fluorescent protein ZsGreen (ZsG). LGR5 mRNA and protein levels were significantly reduced in cells expressing shLGR5, compared to cells expressing shC (Fig. 6A-C). NSCs were then transduced with retrovirus expressing shC or shLGR5, and after 24 hours were co-cultured with astrocytes. Proliferation in transduced NSCs (ZsG+) was assessed by Ki67 staining (Fig. 6D). The percentage of ZsG+ cells positive for Ki67 was significantly reduced in LGR5-deficient NSCs compared to control cells expressing shC (Fig. 6E).

**Figure 6.**
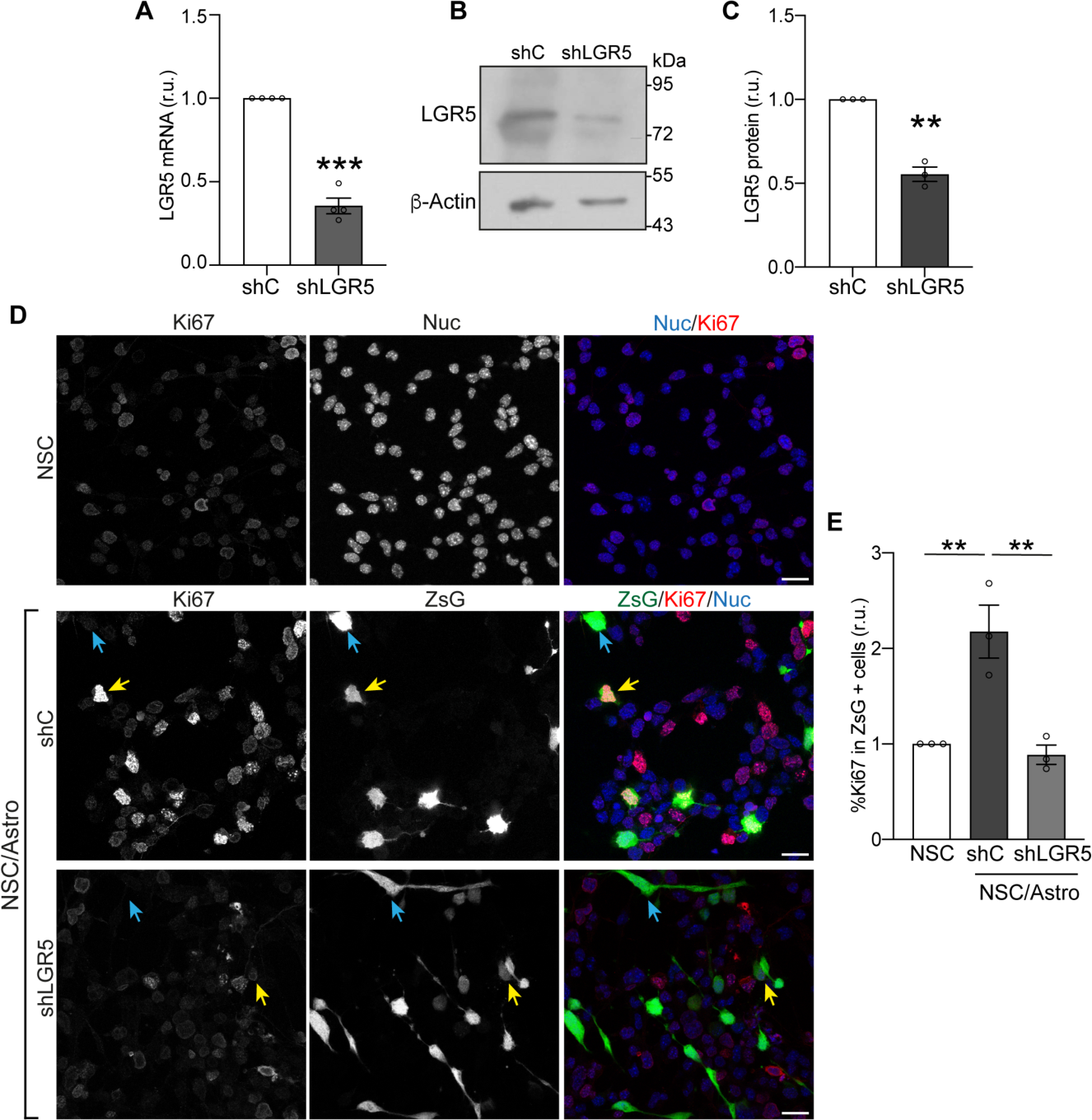
LGR5-deficient hippocampal NSCs have a reduced proliferative response to co-culture with dentate gyrus astrocytes. (A) RT-qPCR from total RNA isolated from NSCs transduced with a retrovirus expressing shRNA targeting LGR5 (shLGR5) or a control shRNA (shC) and the reporter protein ZsG. LGR5 mRNA was normalized to β-actin mRNA and expressed relative to shC. (B) Immunoblot of total protein extracts of NSCs 72 hours after transduction with the retroviruses. (C) Densitometric analysis of the immunoblots. LGR5 protein levels were normalized to β-actin protein levels and expressed relative to the shC condition. (D) Representative immunofluorescence staining of Ki67 in shRNA-transduced cells (ZsG+) in hippocampal NSCs with or without co-culture with astrocytes for 24 hours. Yellow arrows indicate ZsG+Ki67+ cells and blue arrows indicate ZsG+Ki67-cells. (E) Percentage of ZsG+ cells positive for Ki67. Bars represent the mean ± SEM. Dots represent independent experiments. One-sample t test and Wilcoxon test (A, C) or Two-way ANOVA and Tukey‘s post-hoc test (E) were used. **p<0.01, ***p<0.001.

Altogether, these findings demonstrate that RSPO2 secreted by astrocytes and the LGR5 receptor expressed in hippocampal NSCs are crucial for astrocyte-induced proliferation of NSCs.

## DISCUSSION

Our evidence suggests that RSPO signaling is part of the mechanisms that control the proliferation of neural precursor cells in the adult dentate niche. During development and in adult tissues, RSPO proteins enhance Wnt/μ-catenin signaling. Secreted RSPOs interact with a LGR receptor and the E3 ubiquitin ligase ZNRF3 or RNF43 and induce the internalization and degradation of FZD receptors (Hao et al. 2012; Koo et al. 2012). This interaction triggers the membrane clearance of E3 ligase, which in the absence of RSPOs, ubiquitinates FZD receptors inducing their internalization and degradation [reviewed in (de Lau et al. 2014; Zebisch and Jones 2015; Nagano 2019)]. By augmenting Wnt activity, RSPOs exert critical functions in adult tissue homeostasis by controlling proliferation and fate commitment of stem cells in several peripheral tissues (Yan et al. 2017; Harnack et al. 2019; Sigal et al. 2019). Moreover, RSPOs have been used to induce tissue regeneration (Zhao et al. 2009; Zhang et al. 2020) and therefore, they have emerged as interesting therapeutic targets to be used in regenerative medicine applications. Our results suggest that RSPO signaling could be an interesting target to promote neurogenesis in physiological and pathological conditions including aging (Ben Abdallah et al. 2010; Knoth et al. 2010; Moreno-Jimenez et al. 2019), and neurodegenerative diseases (Moreno-Jimenez et al. 2019; Tobin et al. 2019; Terreros-Roncal et al. 2021; Zhou et al. 2022; Cao et al. 2023).

We determined that RSPO1-3 and their LGR4-6 receptors are expressed in the adult mouse dentate gyrus. In agreement with *in situ* hybridization data (Lein et al. 2007), RSPO4 was not detected by RT-qPCR, so this protein was not evaluated in the following experiments. Moreover, RSPO4 is the least active RSPO protein, as determined in cell-based Wnt signaling assays (Kim et al. 2008). Interestingly, the differential distribution of RSPO1-3 determined by immunostaining in hippocampal sections suggests that these ligands might play different roles in the hippocampus. Particularly, RSPOs expressed in the CA3 region might control the reactivation of Wnt signaling reported in mature neurons, as previously suggested (Heppt et al. 2020). Distinct patterns of expression (Nam et al. 2007), as well as divergent roles (Nagano 2019; Ter Steege and Bakker 2021), have been described for RSPOs at different developmental stages.

Furthermore, we found that LGR4-6 receptors are expressed in the dentate gyrus. LGRs have been identified as markers of stem cells in adult tissues (Barker et al. 2007; Jaks et al. 2008; Barker et al. 2010; Mustata et al. 2011; de Visser et al. 2012; Wang et al. 2013; Lehoczky and Tabin 2015; Blaas et al. 2016); however, LGR5 is also expressed in postmitotic cerebellar granule neurons during postnatal development (Miller et al. 2014) and postmitotic neurons in the adult olfactory bulb (Yu et al. 2017), and LGR4 is expressed in Purkinje cells of the cerebellum (Van Schoore et al. 2005) We found that LGR4-5 are expressed in proliferative hippocampal NSCs, suggesting that these cells are sensitive to RSPO proteins. Accordingly, treatment with recombinant RSPO1-3 significantly induced the proliferation of NSCs. RSPOs have previously been shown to be required for proliferation of intestinal stem cells (Kim et al. 2005; Chaves-Perez et al. 2022); RSPO overexpression induces proliferation of these cells, while RSPO inhibition induces lineage commitment and differentiation (Yan et al. 2017). Also, RSPOs increase proliferation of stomach cells (Sigal et al. 2017), adult hair follicle dermal stem cells (Hagner et al. 2020), and lung epithelial stem/progenitor cells (Raslan et al. 2022).

Interestingly, we found that the expression of LGR4-5 receptors decreased upon NSC differentiation, suggesting that RSPO/LGR signaling is reduced after fate commitment. As previously reported, the activity of Wnt/β-catenin signaling also drops upon differentiation. In the dentate gyrus, Wnt/β-catenin activity declined in committed neural progenitors (Garbe and Ring 2012; Heppt et al. 2020; Austin et al. 2021), and in cultured hippocampal NSCs, the Wnt3a-induced TCF/LEF response was gradually downregulated throughout the first 2– 3 days of differentiation (Schafer et al. 2015). Our results suggest that the downregulation of LGR4-5 expression upon NSC differentiation may contribute to the attenuation of the canonical Wnt/β-catenin signaling pathway observed in neural committed progenitors.

Considering that the action of RSPOs involves augmenting the sensitivity of cells to Wnt ligands (Kim et al. 2008), rRSPOs likely increased proliferation by amplifying basal activity of endogenous Wnt ligands expressed in adult hippocampal NSCs (Wexler et al. 2009). In response to rRSPOs treatment, we found increased mRNA levels of the Wnt target gene CyclinD1. Interestingly, we did not observe a potentiation of this effect on CyclinD1 expression or NSCs proliferation by co-treatment with rRSPOs plus rWnt3a. However, treatment with rWnt3a or co-treatment with rRSPOs plus rWnt3a did induce the expression of Axin2 and RNF43, which was not observed by treatment with rRSPOs alone. Both Axin2 and RNF43, are negative regulators of the Wnt/β-catenin signaling, acting at different levels of the Wnt pathway. Axin2 induces the degradation of β-catenin by functional interaction with APC and GSK-3β (Clevers and Nusse 2012) while RNF43 promotes the membrane clearance of FZD receptors (Koo et al. 2012). Mutations in these proteins result in aberrant activation of the Wnt signaling pathway and are found in a variety of human cancers (Liu et al. 2000; Lustig et al. 2002; Hao et al. 2016). Our findings suggest that there is a tight control of Wnt/β-catenin activity in proliferative NSCs. On the one hand, positive signals allow the proliferation of NSCs, while negative feedback mechanisms that limit Wnt activity prevent excessive proliferation. In this context, our results suggest that low levels of Wnt activity induced by treatment with rRSPOs alone induce proliferation, while higher levels of Wnt activity induced by treatment with rWnt3a plus rRSPOs activate the negative feedback, thus blocking further proliferation. This agrees with previous findings in which stimulation of Wnt/β-catenin signaling with the GSK3β inhibitor CHIR99021 reduced the proliferative capacity of cultured hippocampal precursor cells while inducing a strong increase in Axin2 expression (Austin et al. 2021).

In addition, it was reported that high stimulation of Wnt/β-catenin signaling promotes the differentiation of hippocampal NSCs while reducing proliferation (Austin et al. 2021). Moreover, elevated and sustained β-catenin activation sequentially promotes the proliferation and subsequent differentiation of adult NSCs (Rosenbloom et al. 2020). We observed that treatment with Wnt3a ± rRSPOs increased the expression of Ngn2, suggesting that this treatment induced the differentiation of NSCs. In this context, our results suggest that lower levels of Wnt activity induced by treatment with rRSPOs alone induced proliferation, while higher levels of Wnt activity induced by treatment with Wnt3a ± rRSPOs promoted differentiation.

In addition to augmenting Wnt/β-catenin signaling, other ways of action have been described for RSPOs including modulation of non-canonical Wnt signaling (Glinka et al. 2011; Ohkawara et al. 2011; Scholz et al. 2016), and the antagonistic effect on BMP signaling (Lee et al. 2020). We cannot exclude these mechanisms are also involved in the effect of rRSPOs on adult hippocampal NSCs.

Astrocytes are important components of the neurogenic niche that provides signaling molecules at the SGZ and also in the ML close to NSC projections (Sultan et al. 2015; Moss et al. 2016; Casse et al. 2018; Schneider et al. 2019; Arredondo et al. 2022). We determined that astrocytes isolated from the dentate gyrus of adult mice express RSPO1-3. We found that no-contact co-cultures of hippocampal NSCs with dentate gyrus astrocytes induced the proliferation of NSCs, which was prevented by: (i) co-incubation with the extracellular domains (Fc-chimera) of LGR4 or LGR5; (ii) knocking down the expression of RSPO2 in astrocytes; and (iii) knocking down the expression of LGR5 in NSCs. We knocked down the expression of RSPO2 since RT-qPCR data suggested this is the highest expressed RSPO in astrocytes (with lower delta CT value). Similarly, LGR5 was selected since RT-qPCR analysis suggested this is the most highly expressed LGR in NSCs; in addition, Fc-LGR5 strongly blocked proliferation induced by astrocytes.

These data strongly support that RSPO2/LGR5 signaling is involved in the previously described astrocytes-induced proliferation of hippocampal NSCs (Song et al. 2002; Lie et al. 2005; Okamoto et al. 2011). Interestingly, RSPO2 knockdown in astrocytes completely prevented the effect of co-culture on proliferation, even though there was an incomplete decrease of RSPO2 levels in the lysate and media. This suggests that there might be a concentration threshold for RSPO2 to induce proliferation. Additionally, treatment with Fc-LGR4-5 did not affect basal NSCs proliferation, when not in co-culture, possibly because the concentration of RSPOs secreted by NSCs is too low to mediate an autocrine effect. Similarly to what we propose for RSPO levels, Wnt concentration thresholds have been reported to be required for Wnt/β-catenin effects, including cell proliferation in the intestinal crypts (Dunn et al. 2016), and lineage commitment in embryonic stem cells (Tsakiridis et al. 2014).

In summary, our results indicate that RSPO proteins are expressed in the adult dentate gyrus, and RSPOs secreted by dentate astrocytes promote proliferation of hippocampal precursor cells in an LGR-dependent manner. These data indicate that RSPO/LGR signaling might be part of the signaling mechanisms of the adult dentate niche.

## MATERIALS AND METHODS

### Animals

All procedures involving animals were performed according to the NIH and ARRIVE guidelines and were conducted with the approval of the Bioethics Committee of Universidad Andrés Bello (006/2019). Adult two-month-old female and male C57/BL6 mice were used for the experiments. Mice had access to water and food ad libitum in a 12:12 hours light/dark cycle.

### RT-qPCR

Total RNA from NSCs, astrocytes, dentate gyrus, or N2a cells was extracted using TRIzol reagent (Life Technologies) and reversely transcribed into complementary DNA (cDNA) using M-MuLV reverse transcriptase (NEB.M0253S, New Englands BioLabs, Ipswich, MA). qPCR was performed using the Brilliant II SYBR Green QPCR master mix (Agilent Technologies, Santa Clara, CA). The primers used are listed in Supplemental Table S1. Gene expression was normalized to the endogenous housekeeping genes β-actin (ΔCt) and to control samples (ΔΔCt). Fold-change in mRNA levels was calculated using 2^−ΔΔCT^ (Rao et al. 2013). For all samples, technical triplicates were carried out for each qPCR run and then averaged, and at least 3 independent experiments were performed.

### Western blot analysis

Total protein extracts from the dentate gyrus, NSCs, or astrocytes were prepared as previously described (Arredondo et al. 2020a). Proteins were resolved in 10% SDS/PAGE, transferred to a PVDF membrane, blocked, and incubated overnight at 4°C with primary antibodies: rabbit anti-DCX (4604, CellSignaling Technology Inc., 1:750), mouse anti-β-actin (A5441, Sigma-Aldrich, 1:10,000), rabbit anti-GFAP (Z0334, Dako, 1:1000), goat anti-RSPO1 (AF3474, R&D Systems, 1:200), rabbit anti-RSPO2 (MB59206616, MyBioSource, 1:500), rabbit anti-RSPO3 (PA5-38052, Invitrogen, 1:250), rabbit anti-LGR4 (PA5-67868, Invitrogen, 1:500), mouse anti-LGR5 (MA5-25644, Invitrogen, 1:2000), and rabbit anti-LGR5 (HPA012530, Atlas Antibodies, 1:500). Peroxidase-conjugated secondary antibodies (Invitrogen) were then used, and detected using the ECL technique (PerkinElmer, Waltham, MA).

### Perfusion, postfixation, and tissue sectioning

Animals were anesthetized using a mixture of 200 mg/kg ketamine and 20 mg/kg xylazine and transcardially perfused with saline serum, followed by 4% paraformaldehyde (PFA, Sigma-Aldrich) in 0.1 M PBS. Brains were removed and postfixed in 4% PFA in PBS for 24 hours at room temperature, and then dehydrated in 30% sucrose. Brains were sectioned on a cryostat (Cryostat MEV, Slee).

### Immunofluorescence staining

Immunostaining was carried out as previously described for tissue sections (Varela-Nallar et al. 2014) or cultured cells (Varela-Nallar et al. 2009; Abbott et al. 2013). The primary antibodies used were: mouse anti-GFAP (MA515086, Thermo Fischer Scientific, 1:8000), mouse anti-S100β (ab4066, Abcam, 1:100), rat anti-BrdU (ab6326, Abcam, 1:300), rat anti-Ki67 (14-5698-82, Invitrogen, 1:250), mouse anti-Nestin (MAB353, Millipore, 1:50), goat anti-Sox2 (2748, Cell Signaling Technology Inc., 1:100), goat anti-DCX (SC-8066, Santa Cruz Biotechnology, 1:250), rabbit anti-DCX (4604, Cell Signaling Technology Inc., 1:750), goat anti-RSPO1 (AF3474, R&D Systems, 1:200), rabbit anti-RSPO2 (HPA024764, Sigma, 1:500), rabbit anti-RSPO3 (PA5-38052, Invitrogen, 1:150), rabbit anti-LGR4 (PA5-67868, Invitrogen, 1:250), and rat anti-LGR5 (MAB8240SP, R&D Systems, 1:50). As secondary antibodies, Alexa- and DyLight-conjugated antibodies were used. NucBlue (R37605, Invitrogen) was used as a nuclear dye. Image acquisition was performed using a Nikon Ti-E Microscope (Nikon, USA) or a Leica TCS SP8 (Leica Microsystems) confocal microscope using a 40x or 60x objective.

### Isolation and culture of progenitors from the adult mouse hippocampus

NSCs were isolated from the hippocampus of 10 female 6-8-week-old C57/BL6 mice and cultured in monolayers, as previously described (Babu et al. 2011), with some modifications as in (Guerra et al. 2022). NSCs were plated at 10,000 cells/cm^2^ in culture plates pretreated with poly-D-lysine (Merck) and laminin (Gibco), as previously described in (Guerra et al. 2022). NSCs were cultured in DMEM/F12 (Gibco), supplemented with B27 (Gibco), 100 U/ml penicillin and 100 μg/ml streptomycin (Gibco), and the growth factors FGF-2 (20 ng/ml, Alomone Labs) and EGF (20 ng/mL, R&D systems). NSC differentiation was induced withdrawal of these growth factors.

For proliferation analysis, NSCs were treated in the presence or absence of the recombinant proteins RSPO1 (7150-RS/CF, R&D Systems), RSPO2 (6946-RS/CF, R&D Systems), or RSPO3 (4120-RS/CF, R&D Systems) for 24 hours, and 5 μM BrdU (Sigma-Aldrich) was added for the last 2 or 4 hours of treatment. Proliferation was quantified as the percentage of BrdU-positive cells out of the total number of cells positive for the nuclear staining. For differentiation analysis, NSCs were treated with recombinant proteins for 4 days in the absence of growth factors. Neuronal differentiation was quantified as the percentage of cells that expressed DCX.

### Isolation and culture of astrocytes from the adult mouse dentate gyrus

Thew extraction of the dentate gyrus of 6-8-week-old mice was performed as previously described (Walker and Kempermann 2014), and then astrocytes were isolated from this tissue as described (Kaech and Banker 2006), with some modifications. Tissues were rinsed in ice-cold 2M glucose solution in PBS, incubated for 5 min at 37°C with a solution of 0.05% trypsin-EDTA and 250 U/ml DNase I (AM2222, Ambion) in glial medium [MEM (Gibco) supplemented with 0.6% glucose, penicillin/streptomycin, and 10% horse serum], and then mechanically dissociated with fire-polished glass Pasteur pipettes of decreasing diameters. After homogenization, cells were centrifuged for 5 min at 1,500 rpm, the supernatant was removed, and cells were resuspended in glial medium and plated in poly-D-lysine/laminin-coated T-25 bottles. Every 3-4 days, the cells were washed with PBS, and the culture medium was replaced. When reaching 80% confluency, cells were shaken at 60 rpm for 16 hours at 37°C with 5% CO_2_ to remove microglia, and after passaging, the cells were seeded at 2.5 x 10^4^ cells/cm^2^ in poly-D-lysine/laminin-coated plates; the medium was replaced every 4 days. All experiments were performed using astrocytes at <20 passages.

Astrocytes were transfected with lentiviral vectors expressing shRNAs (see the “Lentivirus and retrovirus production” section below) using Lipofectamine 3000 reagent (Invitrogen), following the manufacturer’s instructions. Twenty-four hours after transfection, cells were rinsed with PBS and cultured for 48 hours, in glial medium without horse serum. The media were harvested and centrifuged at 1,500 g for 5 min, and the supernatant was passed through a 0.22 µm filter and precipitated with methanol/chloroform in a ratio of 1:1:0.25. Total cell lysates were prepared as described in the Western blot analysis section above.

### Hippocampal NSCs/Astrocyte sandwich co-culture system

Co-cultures were obtained as previously described (Song et al. 2002; Kaech and Banker 2006), including the following modifications. Briefly, astrocytes were seeded in poly-D-lysine/laminin-coated culture plates at 2.5 x 10^4^ cells per cm^2^ for 4 days. In parallel, hippocampal NSCs were seeded onto poly-D-lysine/laminin-coated glass coverslips in 24-well culture plates at 2 x 10^4^ cells per cm^2^. After 24 hours, NSCs were mounted on 3D discs positioned on the astrocyte monolayer that had been previously cultured in NSC medium for 24 hours. The co-culture discs were 3D printed using dimensions to position a 12 mm coverslip with NSCs 2 mm above the astrocyte cultures (Supplemental Fig S3A).

For the proliferation experiments, co-cultures were maintained for 24 hours in the control condition or in the presence of the chimera proteins Fc-LGR4 (8077-GP, R&D Systems), Fc-LGR5 (8078-GP, R&D Systems), and Fc-Control (110-HG, R&D Systems) for 24 hours, and 10 μM BrdU was added for the last 4 hours of treatment. Proliferation was quantified as the percentage of BrdU-positive cells. For RSPO2 knockdown experiments, astrocytes were transfected with lentiviral vectors expressing shRNAs (see “Lentivirus and retrovirus production” section below) using Lipofectamine 3000 reagent (Invitrogen) following the manufacturer’s instructions. Twenty-four hours post-transfection, the glial medium was replaced by NSCs medium, and after 24 hours, the co-cultures were mounted in the presence of growth factors. After another 24 hours, NSCs were fixed, and proliferation was quantified as the percentage of Ki67-positive cells.

For LGR5 knockdown experiments, NSCs were transduced for 48 hours with shRNA-expressing retroviruses (see “Lentivirus and retrovirus production” section below) and subsequently co-cultured with or without astrocytes for 24 hours in the presence of growth factors. Proliferation was quantified as the percentage of Ki67-positive cells.

### Lentivirus and retrovirus production

To knockdown RSPO2 and LGR5, inverted self-complementary hairpin DNA oligonucleotides encoding short-hairpin RNAs (shRNA) against mouse RSPO2 (shR1 and shR2) or LGR5 (shLGR5) were chemically synthesized. Oligos used to construct the shRNAs were: shR1: 5’-TG*CGAGCTAGTTATGTATCAAAT*TTCAAGAGAATTTGATACATAACTAGCTCGCTT TTTTC-3’; shR2: 5’-TG*AGCGAGCTAGTTATGTATCAA*TTCAAGAGATTGATACATAACTAGCTCGCTC-TTTTTTC-3’; shLGR5: 5’-GA*TCCGGCGAGTCTGCTGTCCATTAAC*TTCAAGAGAGTTAATGGACAGCAGACTC GCTTTTTTACGCGTG-3’. Target mRNA sequences are shown in italic. As a control, shRNA targeting luciferase mRNA (Varas-Godoy et al. 2018) was used. RSPO2 shRNAs were cloned into the pLentiLox 3.7 lentiviral vector (PLL3.7, AddGene), which co-expresses green fluorescent protein (GFP), and shLGR5 was cloned into the pSiren-RetroQ-ZsGreen vector (Clontech). For lentivirus preparation, 1×10^6^ HEK293T cells were seeded per 100 mm dish, once the cells reached 90% confluence (∼3 days), cells were co-transfected with 14.4 μg pLentiLox vector containing the sequence of each shRNA, 9.2 μg pMDLg/pRRE vector encoding the gag and pol enzymes, 4.9 μg pVSV-G vector required for capsid formation, and 3.6 µg pRSV-Rev vector using 65 µL of PEImax (18 mM polyethylenimine, pH 7, Polysciences, Inc.). After 4 hours, the medium was replaced by fresh DMEM medium supplemented with 1% fetal bovine serum (FBS, Biowest) without antibiotics. The lentivirus-containing supernatants were collected 72 hours post-transfection, and centrifuged twice at 2,000 g for 30 min at 4°C on AMICON filters (Ultra-15 Centrifugal Filter Unit). The concentrated viruses were aliquoted and stored at −80°C.

For retrovirus production 1×10^6^ HEK293T cells were seeded in 100 mm culture dishes, and once the cells reached 90% confluence (∼3 days), they were co-transfected with 220 μL Optimem, 14.4 μg silencing vector, 9.2 μg pMDL g/p, 4.9 μg VSV-G, and 65 μL PEImax. The retroviral transfection mix was incubated for 10 min at room temperature, added dropwise onto the HEK293T cell medium, and left at 37°C with 5% CO_2_. After 4 hours, the medium was replaced by DMEM medium supplemented with 1% FBS. At 72 hours post-transfection, the supernatants were harvested and centrifuged at 2,000 g for 5 min at 4°C. Next, the supernatants were filtered using 0.22 μm cellulose acetate filters and then underwent two rounds of ultracentrifugation: 19,000 rpm for 2 hours, and then at 21,900 rpm for 2 hours. The pellet was resuspended in PBS, aliquoted, and stored at −80°C.

### Statistical analysis

Statistical analyses were performed using Prism 10 software (GraphPad Software Inc.). Data normality was checked using the Shapiro-Wilk test. One sample t-test or Wilcoxon test was used for fold-change analyses. A two-way ANOVA followed by a Tukey’s or Dunnett’s multiple comparison test was used in experiments with two variables. *p*<0.05 was considered statistically significant. Data represent the mean ± SEM.

## Acknowledgments

This work was supported by grants from ANID/FONDECYT (1190461 and 1230454, to LVN), Nucleo UNAB (DI-4-17/N, to LVN), ANID 21210618 to DVB, and ANID/FONDECYT postdoctoral grant (3230100, to SBA).

## Author contributions

DV-B: conception and design, retrovirus production, acquisition, analysis, and interpretation of data. MDM: conception and design, and acquisition of data. MG: lentivirus production, data acquisition and analysis of co-culture experiments with RSPO2-deficient astrocytes. SBA: acquisition and analysis of data from RSPO2 knockdown. SHS: Lentivirus production and validation of RSPO2 knockdown. NM-V: design and production of the sandwich co-culture system. FJB: shRNAs design (RSPO2 and LGR5), lentivirus production, analysis, and interpretation of data. LV-N: conception and design, assembly and interpretation of data, manuscript writing, and financial support. All authors read and approved the final version of the paper.

